# Library-free BoxCarDIA solves the missing value problem in label-free quantitative proteomics

**DOI:** 10.1101/2020.11.07.372276

**Authors:** Devang Mehta, Sabine Scandola, R. Glen Uhrig

## Abstract

The last decade has seen significant advances in the application of quantitative mass spectrometry-based proteomics technologies to tackle important questions in plant biology. The current standard for quantitative proteomics in plants is the use of data-dependent acquisition (DDA) analysis with or without the use of chemical labels. However, the DDA approach preferentially measures higher abundant proteins, and often requires data imputation due to quantification inconsistency between samples. In this study we systematically benchmarked a recently developed library-free data-independent acquisition (directDIA) method against a state-of-the-art DDA label-free quantitative proteomics workflow for plants. We next developed a novel acquisition approach combining MS^1^-level BoxCar acquisition with MS^2^-level directDIA analysis that we call BoxCarDIA. DirectDIA achieves a 33% increase in protein quantification over traditional DDA, and BoxCarDIA a further 8%, without any changes in instrumentation, offline fractionation, or increases in mass-spectrometer acquisition time. BoxCarDIA, especially, offers wholly reproducible quantification of proteins between replicate injections, thereby addressing the long-standing missing-value problem in label-free quantitative proteomics. Further, we find that the gains in dynamic range sampling by directDIA and BoxCarDIA translate to deeper quantification of key, low abundant, functional protein classes (e.g., protein kinases and transcription factors) that are underrepresented in data acquired using DDA. We applied these methods to perform a quantitative proteomic comparison of dark and light grown Arabidopsis cell cultures, providing a critical resource for future plant interactome studies. Our results establish BoxCarDIA as the new method of choice in quantitative proteomics using Orbitrap-type mass-spectrometers, particularly for proteomes with large dynamic range such as that of plants.

## Introduction

The last decade has seen significant advances in the application of quantitative mass spectrometry-based proteomics technologies to tackle important questions in plant biology. This has included the use of both label-based and label-free quantitative liquid-chromatography mass spectrometry (LC-MS) strategies in model^1,2^ and non-model plants^3^. While chemical labelling-based workflows (e.g. iTRAQ and TMT) are generally considered to possess high quantitative accuracy, they nonetheless suffer from ratio distortion and sample interference issues^4,5^, while being less cost-effective and offering less throughput than label-free approaches. Consequently, label free quantification (LFQ) has been widely used in comparative quantitative experiments profiling the native^6^ and post-translationally modified (PTM-ome)^7,8^ proteomes of plants. However, LFQ shotgun proteomics studies in plants have so far, almost universally, used data-dependent acquisition (DDA) for tandem MS (MS/MS) analysis.

In a typical DDA workflow, elution groups of digested peptide ions (precursor ions) are first analysed at the MS^1^ level using a high-resolution mass analyser (such as modern Orbitrap devices). Subsequently, selected precursor ions are isolated and fragmented, generating MS^2^ spectra that deduce the sequence of the precursor peptide (**Figure 1 a**). For each MS1 scan usually around 10-12 MS^2^ scans are performed after which the instrument cycles to the next MS^1^ scan and the cycle repeats. While this “TopN”selection approach enables identification of precursors spanning the entire mass range, the fragmentation of semi-stochastically selected precursor ions (generally, more intense ions) limits the reproducibility of individual DDA runs, results in missing values between replicate runs, and biases quantitation toward more abundant peptides^9^. This is particularly disadvantageous for label-free workflows and samples with a high dynamic range proteomes, such as human plasma and photosynthetic tissue.

**Figure 1:**
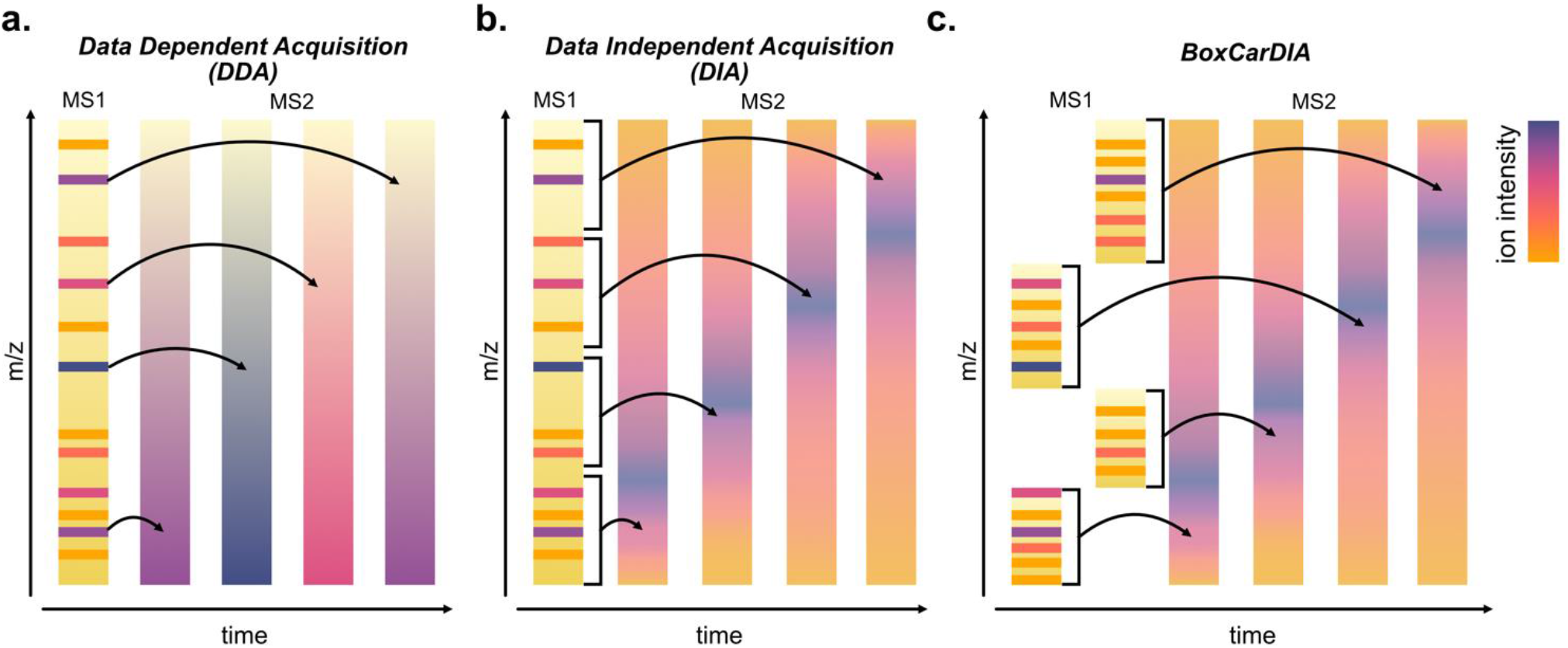
Data acquisition schemes in liquid chromatography mass-spectrometry proteomics. Mass-spectrometry workflows use different acquisition schemes to analyse peptide ions (MS1 analysis) and their corresponding fragment ions (MS2 analysis) to derive peptide sequence information. This study compares three main acquisition methods. **(a.)** Conventional data-dependent acquisition (DDA) involves performing a single MS1 analysis scan followed by the selection of the most intense peptides for fragmentation an MS2 analysis. **(b.)** Data-independent acquisition (DIA) schemes select windows of MS1-analysed peptide ions for fragmentation together, essentially performing MS2 analysis of all MS1-analysed ions rather than on only selected ones. **(c.)** BoxCarDIA seeks to increase the resolution of the MS1 analysis by partitioning it into sequentially analysed sets of boxes, each of which can then be analysed using the DIA approach. This permits better profiling at both the MS1 and MS2 level.

In order to address these limitations, several data-independent acquisition (DIA) workflows have been pioneered, famously exemplified by Sequential Window Acquisition of All Theoretical Mass Spectra (SWATH-MS)^10,11^. In DIA workflows, specific, often overlapping, m/z windows spanning a defined mass range are used to sub-select groups of precursors for fragmentation and MS^2^ analysis. As a result, complete fragmentation of all precursors in that window follows MS^1^ scans resulting in a more reproducible and complete analysis. A major disadvantage of DIA workflows, however, is that each MS^2^ scan contains multiplexed spectra from several precursor ions making accurate identification of peptides difficult. Traditionally, this has been addressed through the use of global or project-specific spectral-libraries obtained from a fractionated, high-resolution DDA survey of all samples—adding to experimental labour and instrumentation analysis time. More recently, alternative approaches have been developed that avoid the use of spectral libraries and instead use “pseudo-spectra” derived from DIA runs that are then searched in a spectrum-centric approach analogous to conventional DDA searches^12–14^. Improvements in such library-free DIA approaches have included the incorporation of high precision indexed Retention Time (iRT) prediction^15^ and the use of deep-learning approaches^16–18^. DirectDIA (an implementation of a library-free DIA method; Biognosys AG) and a hybrid (directDIA in combination with library-based DIA) approach has been recently used to quantify more than 10,000 proteins in human tissue^19^ and reproducibly identify >10,000 phosphosites across hundreds of human retinal pigment epithelial-1 cell line samples^20^.

While DIA addresses the stochasticity of precursor selection for fragmentation, it does not solve the problem of incomplete MS^1^ analysis due to the limited charge capacity of C-traps that lie upstream of Orbitraps. In effect this means that modern Orbitrap mass-spectrometers only analyse <1% of available ions at the MS^1^ level^21^. In 2018, Meier et al., described a novel acquisition scheme called BoxCar where multiple overlapping sets of narrow m/z segments are scanned at the MS^1^ level followed by conventional DDA-type MS^2^ analysis^21^. It is thus reasonable to speculate that combining the power of BoxCar to produce higher-resolution MS^1^ data with library-free DIA-type MS^2^ analysis (BoxCarDIA) may provide greater quantitative depth and range for shotgun proteomics.

DirectDIA combines the advantages of DIA for reproducible quantification of proteins in complex mixtures with high dynamic range, with the ease of use of earlier DDA methodologies. BoxCarDIA can improve MS^1^ resolution and dynamic range, while addressing the limitations of DDA-type precursor fragmentation. Hence, a systematic comparison of these different technologies for LFQ proteomics is essential to define best practice in plant proteomics. In order to execute this analysis, we compared the proteomes of light- and dark-grown Arabidopsis suspension cells generated with DDA, directDIA and BoxCarDIA acquisition schemes. Arabidopsis suspension cells are a long-established platform for plant biochemistry and have recently seen a resurgence in popularity due to their utility in facilitating protein interactomic experimentation using technologies such as tandem affinity purification-mass spectrometry^22–26^, nucleic acid crosslinking^27^, and proximity labelling (e.g. TurboID)^28^. Despite this, no existing resource profiling the basal differences in proteomes of Arabidopsis cells grown in light or dark exists—and as we will demonstrate here, is a fundamental requirement to determine the choice of growth conditions to maximize the utility of protein interactomic experiments and targeted proteomic assays in this system.

## Results & Discussion

### Library-free DIA approaches outperform DDA in quantitative depth

We performed total protein extraction under denaturing conditions from Arabidopsis (cv. Ler) suspension cells grown for five days in either constant light or dark. Trypsin digestion of the extracted proteome was performed using an automated sample preparation protocol, with 1ug of digested peptide subsequently analysed using an Orbitrap Fusion Lumos mass spectrometer operated in either DDA, DIA, or BoxCarDIA acquisition modes over 120-minute gradients. Two separate experiments were performed using independent digests of the extracted Arabidopsis proteins. The first to compare DDA and directDIA, and the second to compare directDIA with BoxCarDIA. Eight injections (4 light & 4 dark) per analysis were carried out. DDA data processing was performed using MaxQuant, while DIA data processing was performed using Spectronaut v14 (Biognosys AG.). For DIA analysis, both hybrid (library+directDIA) and directDIA analysis was performed. The hybrid analysis was performed by first creating a spectral library from DDA raw files using the Pulsar search engine implemented in Spectronaut, followed by a peptide centric DIA analysis with DIA raw output files. DirectDIA and BoxCarDIA analysis was performed directly on raw DIA files as implemented in Spectronaut. For BoxCarDIA analysis, the crucial box size parameter was specified using a custom script (see Methods) that designs boxes with equal number of peptide ions using spectral data from a prior directDIA run. The entire workflow is depicted in **Figure 2**. Both hybrid DIA and directDIA analysis substantially outperformed DDA analysis with an average of 65,351; 57,503; and 42,489 peptide-spectrum matches (precursors) quantified across all 8 samples for each analysis, respectively. Hybrid DIA and directDIA also displayed similar gains over DDA in terms of quantified peptides and protein groups (**Figure 2**). While hybrid DIA analysis performed marginally better than directDIA, further analysis was performed with the results of only directDIA and DDA analyses in order to compare methods that use an equivalent amount of input data, comparable instrumentation time and relatively comparable data analysis workflows. We also found improvements in quantifying precursors, peptides, and protein groups using BoxCarDIA as compared to directDIA. Overall, our results suggest that library-free BoxCarDIA can increase quantitative depth by as much as 40% over conventional DDA methods with no increase in analysis time or change in instrumentation.

**Figure 2:**
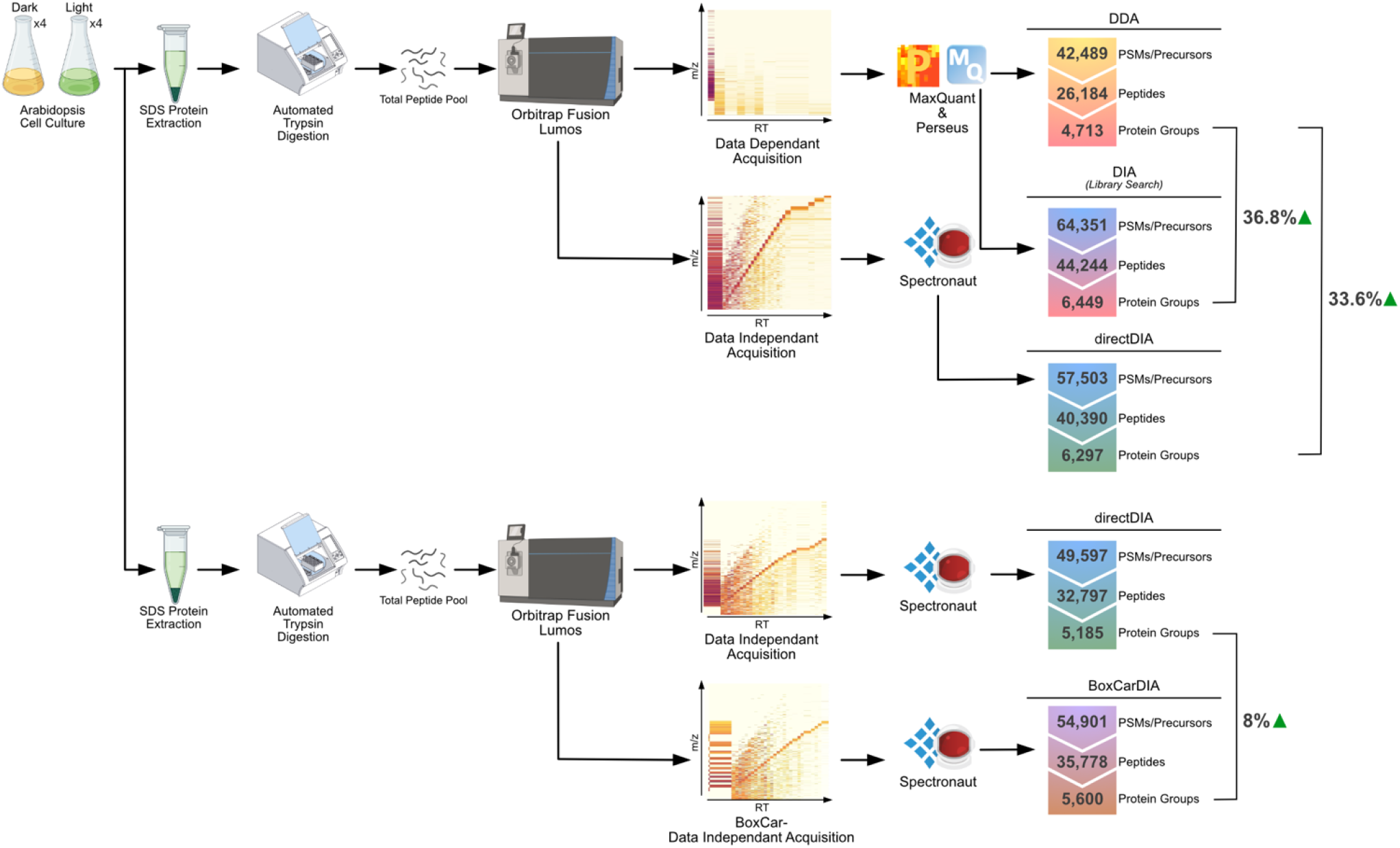
Experimental workflow and summary results. Total protein was isolated from light and dark grown Arabidopsis cells under denaturing conditions for use in two experiments. In the first experiment, peptides were digested with trypsin, desalted and subjected to LC-MS/MS using two different acquisition modes. Ion maps showing a single MS1 scan and subsequent MS2 scans are presented to illustrate differences in acquisition schemes. Raw data was analyzed using MaxQuant & Perseus for data-dependent acquisition (DDA) analysis and Spectronaut for data-independent acquisition (DIA) analysis using spectral libraries created from both acquisitions. Spectronaut was also used for directDIA analysis without the use of spectral libraries. A second experiment involved analyzing independent digests of the same protein extracts followed by the same general analysis pipeline, in order to directly compare directDIA and library-free BoxCarDIA acquisition modes. Counts of FDR-filtered (0.01) peptide spectrum matches (PSMs)/precursors, peptides, and protein groups for each analysis type are shown. Percentage values for increases in protein group quantifications are shown alongside each analysis.

Next, we undertook a series of data analyses to compare the completeness, quality, and distribution of protein group-level quantification of the DDA and directDIA analyses. In order to compare quantification results across the different analysis types, raw intensity values for each sample were log_2_ transformed, median-normalized (per sample), and then averaged for each condition to produce a normalized protein abundance value. For DDA analysis, the number of proteins quantified was determined by first filtering for proteins with valid quantification values in at least 3 of 4 replicates in either condition (light or dark) and then imputing missing values using MaxQuant with standard parameters^29,30^. For directDIA and BoxCarDIA analyses, quantified proteins were defined as those passing standard Q-value filtering in Spectronaut. In total, DDA analysis resulted in the quantification of 4,837 proteins (both conditions) and directDIA analysis quantified 6,526 proteins (light) and 6,454 proteins (dark) (**Supplementary Tables 1-3**). Upon comparing the quantified proteins between both methods, we found that 4,599 proteins were quantified by both techniques, 1,934 were quantified only by directDIA and 235 proteins were exclusively DDA-quantified, for light-grown cells (**Figure 3a**). A correlation plot of normalized quantification values for the 4,599 common proteins showed a moderate correlation between DDA and directDIA quantification (Spearman’s R = 0.773) (**Figure 3a**). Examining the frequency distribution of proteins quantified in light-grown cells, by both methods, revealed that the DDA results were substantially skewed towards higher abundant proteins compared to directDIA (**Figure 3b**). In order to investigate the overlap of quantified proteins between directDIA and DDA at extreme protein abundances, we sub-selected the 2%, 5%, 95% and 98% percentile of the combined quantification distribution and constructed UpSet plots^31^ for these datasets. This analysis revealed that directDIA quantifies hundreds of more proteins at the lower extremes but is only marginally less effective than DDA at the upper extremes of the protein abundance distribution (**Figure 3c**). These results were similarly replicated for dark-grown cells, suggesting that this is a universal feature of the two acquisition methods, irrespective of sample treatment or type (**Figure 3 d-f**). In order to assess if this difference in quantification ability is specific to plant cells (that have a high dynamic range of protein levels), we further analysed a commercial HeLa cell digest standard using the same mass spectrometry and chromatography settings, with quadruplicate injections per analysis type. Analysing the HeLa quantification results (**Supplementary Tables 4 & 5**) showed a similarly uniform quantification across a wide range by directDIA and a slightly better, but still skewed, performance by DDA compared to Arabidopsis cells (**Figure S1 a & b**). Comparing the quantification values for HeLa proteins acquired by directDIA and DDA showed a stronger correlation than for Arabidopsis (Spearman’s R=0.886). Indeed, correlations between quantification values for lower abundant proteins (defined here as proteins below the median quant value), were much lower than for the overall dataset in both species, and yet slightly stronger in the case of HeLa proteins (**Figure S1 c-e**).

**Figure 3:**
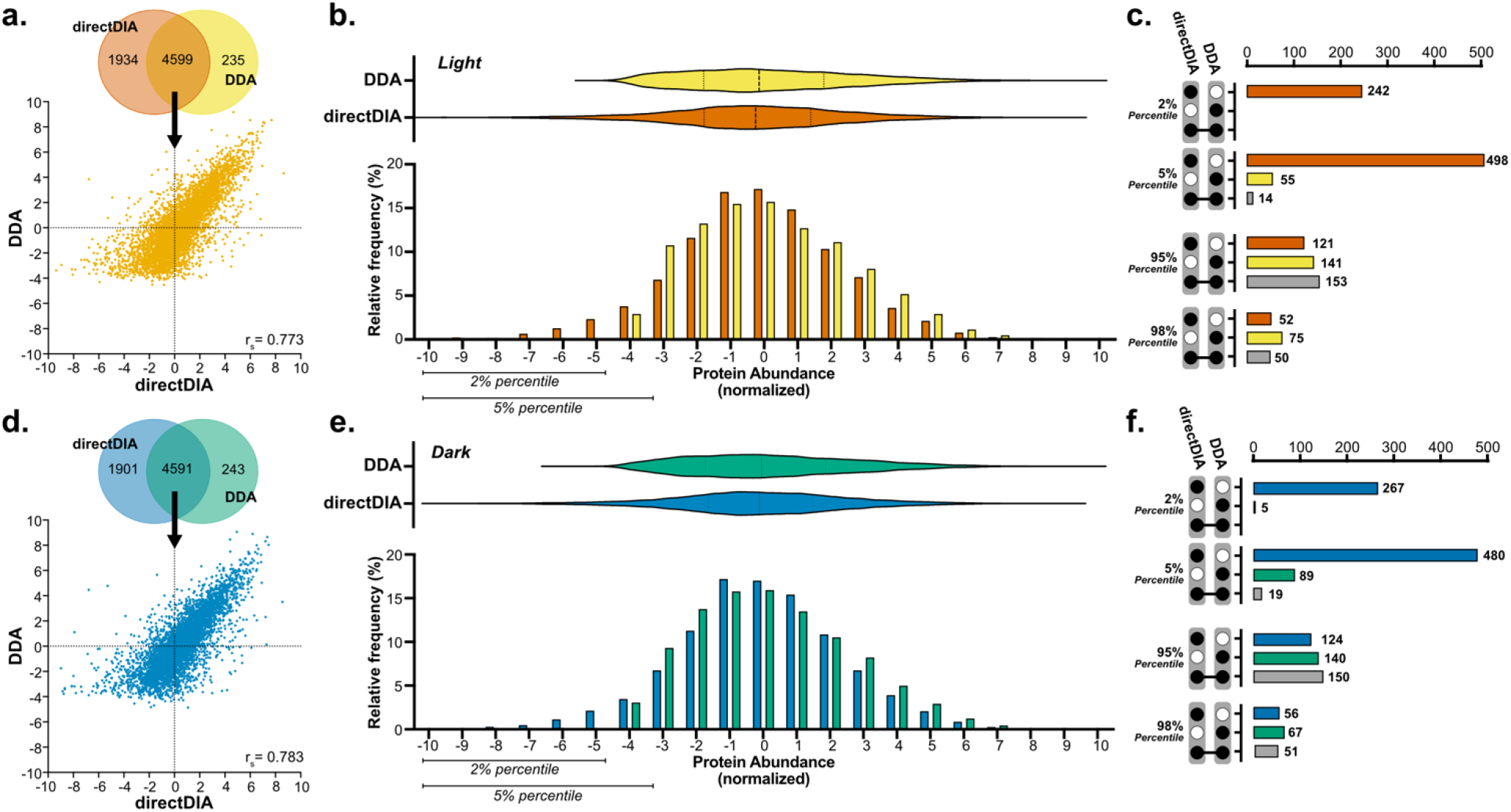
Comparison of protein quantification results using DDA and direct DIA analysis for (a.-c.) light grown and (d.-f.) dark grown Arabidopsis cells. **(a.)** & **(d.)** Venn diagram of protein groups quantified with direct DIA and DDA and scatter plot of protein groups quantified by both methods. r_s_: Spearman’s correlation coefficient. **(b.)** & **(e.)** Frequency distribution of normalized protein abundances for DDA and direct DIA analysis and corresponding violin plots with median and quartile lines marked. **(c.)** & **(f.)** Upset plots depicting intersections in protein groups quantified by DDA and direct DIA at either extremes of the abundance distribution.

We next performed similar comparative analyses for an independent experiment comparing directDIA and BoxCarDIA approaches (**Figure 4**). In this experiment, BoxCarDIA resulted in the quantification of 5,806 (light) and 5,791 (dark) proteins compared to 5,377 (light) and 5,354 (dark) using directDIA (**Supplementary Tables 6 & 7).** The relative abundance of proteins quantified in both analyses correlated to a large degree (Spearman’s r ~ 0.92; **Figure 4 a & d**), much more than the correlation between directDIA and DDA analyses (**Figure 3 a & d**). The frequency distributions of normalised abundances of proteins quantified by both directDIA and BoxCarDIA showed that BoxCarDIA is better able to quantify both high- and low-abundant proteins, for both light and dark grown cells (**Figure 4 b & e**). This is clearly evident upon UpSet plot visualization of the overlap between the two techniques at the extremes of the protein abundance distributions (**Figure 4 c & f**).

**Figure 4:**
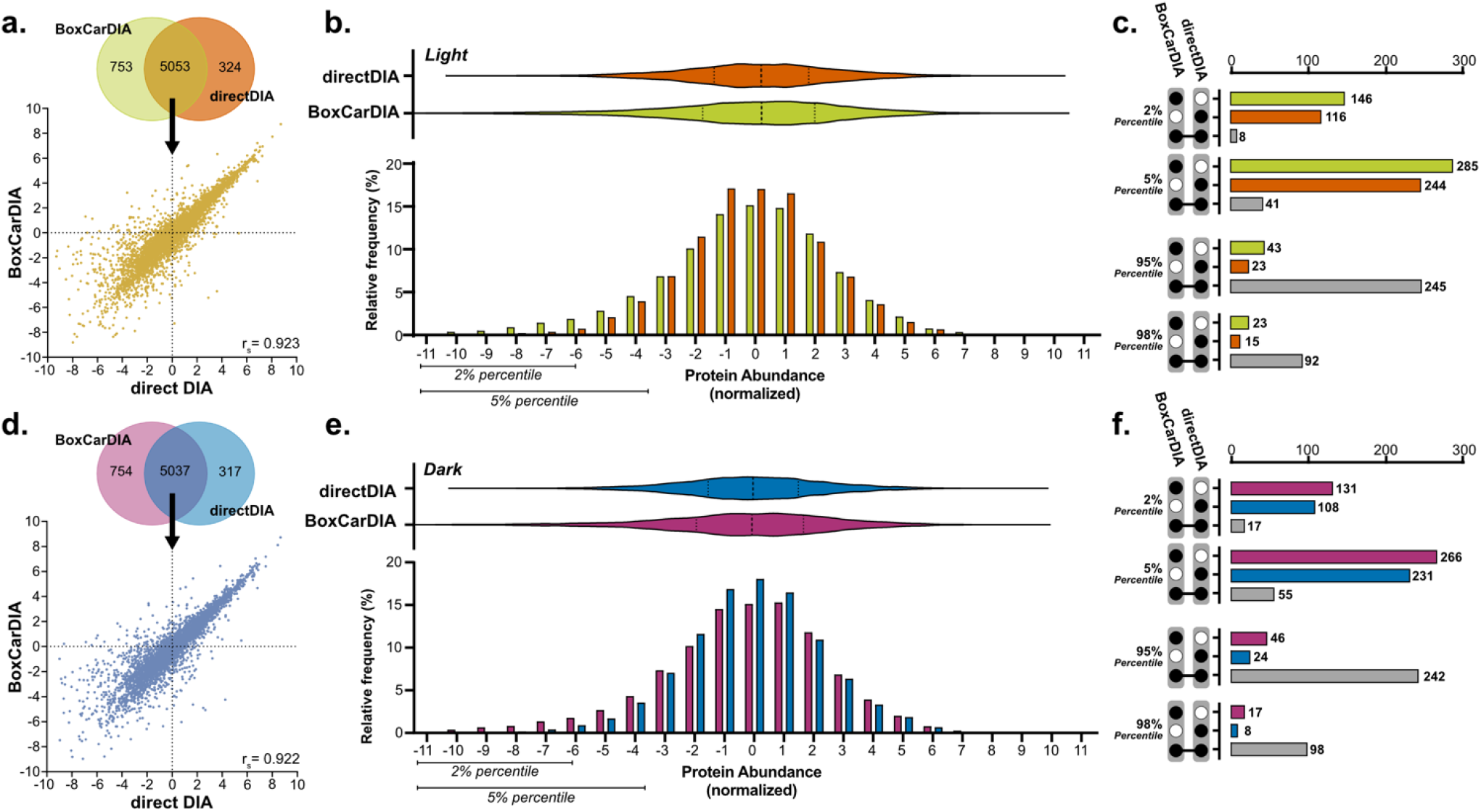
Comparison of protein quantification results using directDIA and BoxCarDIA analysis for (a.-c.) light grown and (d.-f.) dark grown Arabidopsis cells. **(a.)** & **(d.)** Venn diagram of protein groups quantified with BoxCarDIA and directDIA, and scatter plot of protein groups quantified by both methods. r_s_: Spearman’s correlation coefficient. **(b.)** & **(e.)** Frequency distribution of normalized protein abundances for directDIA and BoxCarDIA analysis and corresponding violin plots with median and quartile lines marked. **(c.)** & **(f.)** Upset plots depicting intersections in protein groups quantified by directDIA and BoxCarDIA at either extreme of the abundance distribution.

### BoxCarDIA and directDIA result in more reproducible quantification of peptides and protein groups

In order to deduce the underlying factors limiting the ability of DDA to quantify low abundant proteins, we next investigated quantification distributions for both DDA and directDIA derived data after various data-filtering steps (**Figure S2**; **Supplementary Tables 8-17**). We found that DDA was indeed able to identify a similar number of proteins as directDIA for both Arabidopsis cells and HeLa digests. Predictably these numbers dropped dramatically upon filtering proteins for only those with valid quantification values across 3 of 4 replicates, with only mild gains realized due to imputation of missing values. In contrast, even upon filtering for valid values across 4 of 4 replicates, directDIA resulted in the quantification of more than 5,400 proteins compared to 3,600 complete quantifications for DDA. Strikingly, quantification distributions remained unchanged regardless of various types of data-filtering for directDIA but were greatly skewed towards high abundance upon filtering for valid values in 3 of 4 replicates in DDA outputs (**Figure S2**). This suggests that the poor quantification of low abundant proteins is related to the presence of missing values in DDA analysis.

This hypothesis was reinforced when we distributed the protein quantification data for directDIA and DDA based on the number of replicates with valid quantification values for each protein (**Figure S3**). Here we found that the overwhelming majority (>95%) of proteins quantified by directDIA had valid values in at least 3 of 4 biological replicates for Arabidopsis cells grown in the light or dark (**Figure S3**). In contrast, only 68% and 74% of proteins were accurately quantified by DDA in 4 of 4 replicates of light and dark grown Arabidopsis cells, respectively. When these distributions were further plotted against the normalized protein quantification values, it became clear that proteins found in a lower number of replicates trended lower in abundance in DDA, while this trend did not hold true for directDIA (**Figure S3**). However, the presence of missing values between biological replicates in both methods may yet be explained by real variation between samples.

To account for this, we performed an additional series of experiments by pooling our eight Arabidopsis digests and performing four replicate injections using DDA, directDIA, and BoxCarDIA, respectively. Using replicates that should have the exact protein content allowed us to measure the reproducibility of each data acquisition approach (**Figure 5**). This analysis found that 99.94% of peptides quantified using BoxCarDIA were found in all four Arabidopsis technical replicates, with the remaining 0.06% found in three of four replicates. DirectDIA resulted in slightly less reproducible results, however, only 58.2% of peptides quantified using DDA were found in all four injections. Further, a striking 12% of peptides were quantified by DDA in only one of four technical replicates (**Figure 5 b**). These differences in quantitative completeness at the peptide-level translate to even greater differences at the level of protein groups (**Figure 5 e-h**). Here, we found that nearly a third of proteins groups quantified using DDA were found in only one of four technical replicates, whereas all proteins were quantified in all four replicates by BoxCarDIA. These results were also replicated using a HeLa cell digest, reinforcing their validity (**Figure S4**).

**Figure 5:**
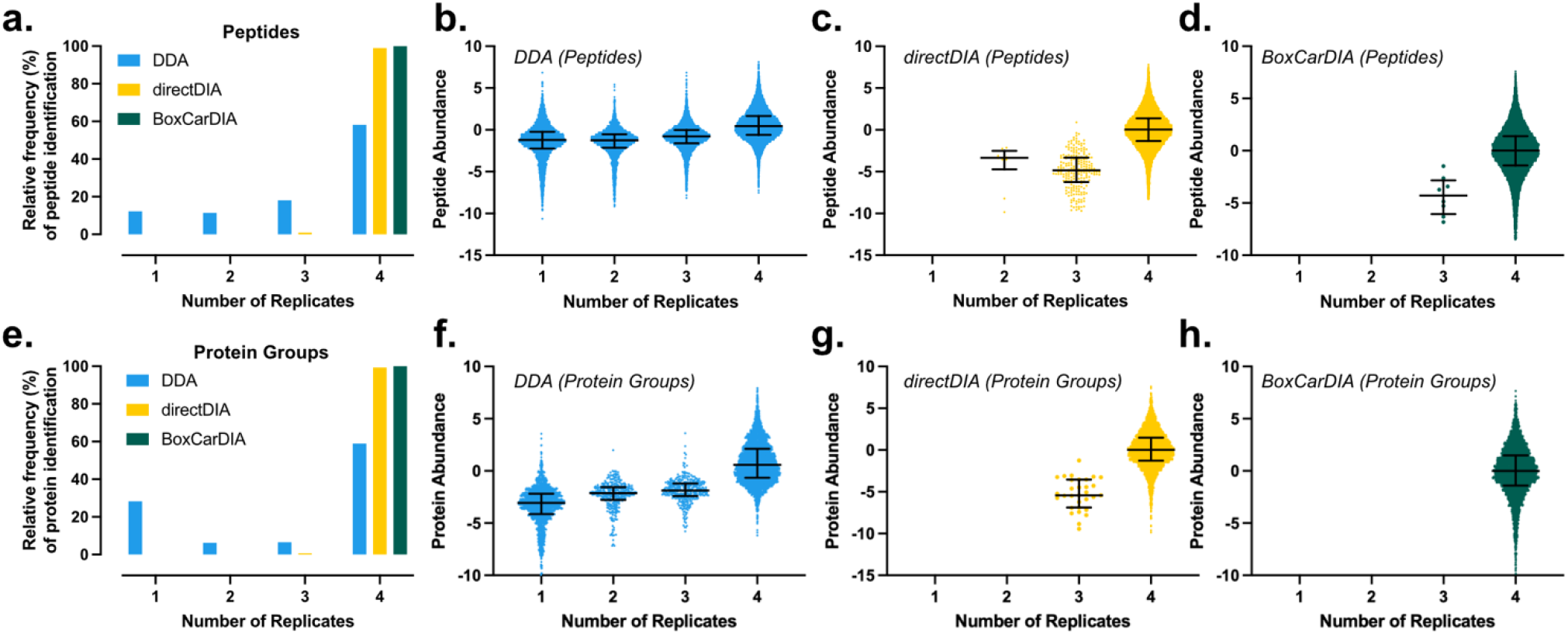
BoxCarDIA can quantify peptides and protein groups consistently between independent technical replicate injections. **(a.)** Histograms of BoxCarDIA, directDIA, or DDA peptide identifications across replicate injections of Arabidopsis cell culture digests. **(b-d.)** Normalized abundances of peptides binned by the number of replicates containing each protein for DDA, directDIA and BoxCarDIA. Bars represent median and interquartile range. **(e-h)** Same as above for protein group identifications.

Like in our analysis of biological replicates, lower abundant proteins tended to be less reproducibly measured across technical replicates in both Arabidopsis and HeLa digests.

Overall, these results reinforce the fact that DDA acquisition results in inconsistent quantification between injections, and that this may in fact obscure real biological variance between samples, especially with regards to lower abundant proteins. Our results also show that the gains in quantitative depth and range provided by better sampling of the ion beam at the MS^1^ level in BoxCarDIA also translate to a complete data matrix, eliminating the long-standing missing-value problem in label-free quantitative proteomics.

### Low abundant protein groups are better represented in BoxCarDIA and directDIA analyses

We next investigated how the better quantitation of lower abundant proteins by directDIA and BoxCarDIA might affect the detection and quantification of biologically functional protein groups in Arabidopsis. We first took advantage of a recently published deep proteome analysis of Arabidopsis tissues^32^ to plot the abundance distributions of all quantified proteins grouped by Gene Ontology categories (**Figure S5**). This allowed us to identify important protein classes with high (glycolysis, translation), medium (proteolysis, carbohydrate metabolism), and low (protein phosphorylation, transcription factors) abundance. We next plotted all proteins detected in our directDIA analysis ranked by abundance, overlayed with the mean abundance of proteins in each of the high, medium, and low abundant classes to verify the classification in our dataset (**Figure 6a**). We then compared the representation of these groups of proteins within the DDA, directDIA, and BoxCarDIA Arabidopsis datasets. Our results show that DDA analysis has a significant over-representation of proteins belonging to high-abundant classes (glycolysis and translation) and a significant underrepresentation of low-abundant proteins involved in protein phosphorylation and transcription (**Figure 6b**). Similarly, BoxCarDIA improves upon directDIA with a significantly enhanced representation of low abundant protein classes (**Figure 6c**). This analysis demonstrates that the better quantification of low abundant proteins by directDIA and BoxCarDIA has consequences on the ability of proteomics studies to measure functionally important regulatory proteins like kinases, phosphatases and transcription factors.

**Figure 6:**
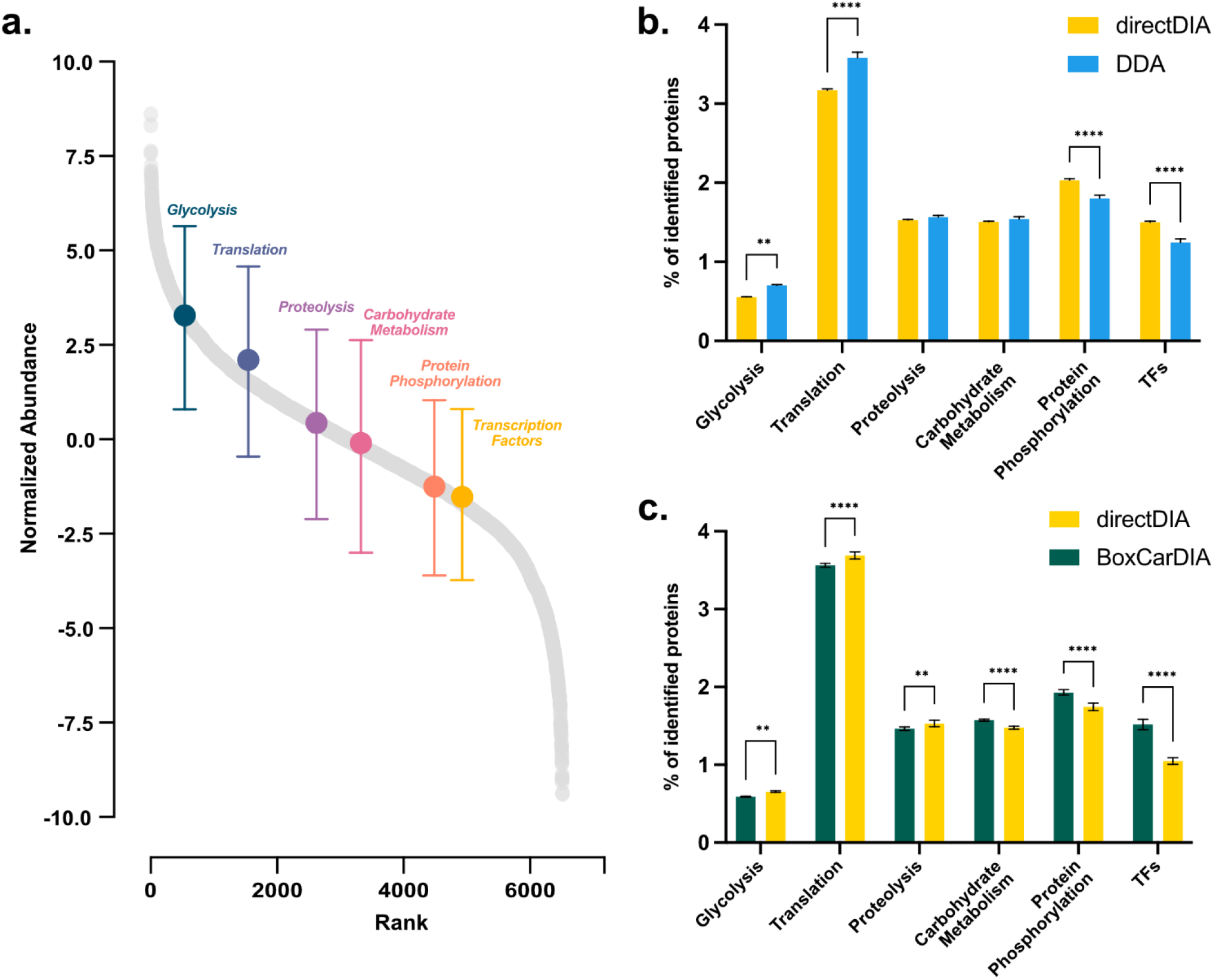
BoxCarDIA and directDIA result in better quantification of low abundant classes of proteins. **(a.)** All proteins quantified by directDIA analysis of Arabidopsis light and dark cell cultures ranked by abundance with the respective high, medium, and low abundant Gene Ontology categories overlayed by mean abundance of component proteins. **(b.)** Percentage of identified proteins in each functional protein group measured by DDA and directDIA analysis, and **(c.)** by directDIA and BoxCarDIA analysis of Arabidopsis light and dark cell cultures. (** p-value<0.01, *** p-value<0.001, **** p-value <0.0001; Šìdàk’s multiple comparisons test)

### A third of all quantified proteins are differentially abundant in light vs. dark grown Arabidopsis cells

Having systematically investigated the advantages and limitations of BoxCarDIA, directDIA and DDA acquisition for LFQ proteomics, we next performed a differential abundance analysis comparing the proteomes of light- and dark-grown cell cultures quantified in our initial directDIA and DDA experiment. We found 2,089 proteins changing significantly in their abundance (Absolute Log2FC > 0.58; q-value <0.05) in our first directDIA analysis and 1,116 proteins changing significantly (Absolute Log2FC > 0.58; q-value <0.05) in DDA analysis. Of these, 710 proteins were found to change significantly in both analyses (**Figure 7a**). In our second experiment, we found 1,639 and 1,920 proteins changing significantly in abundance between light and dark grown cells in our directDIA and BoxCarDIA analyses, respectively (**Figure 7b**).

**Figure 7:**
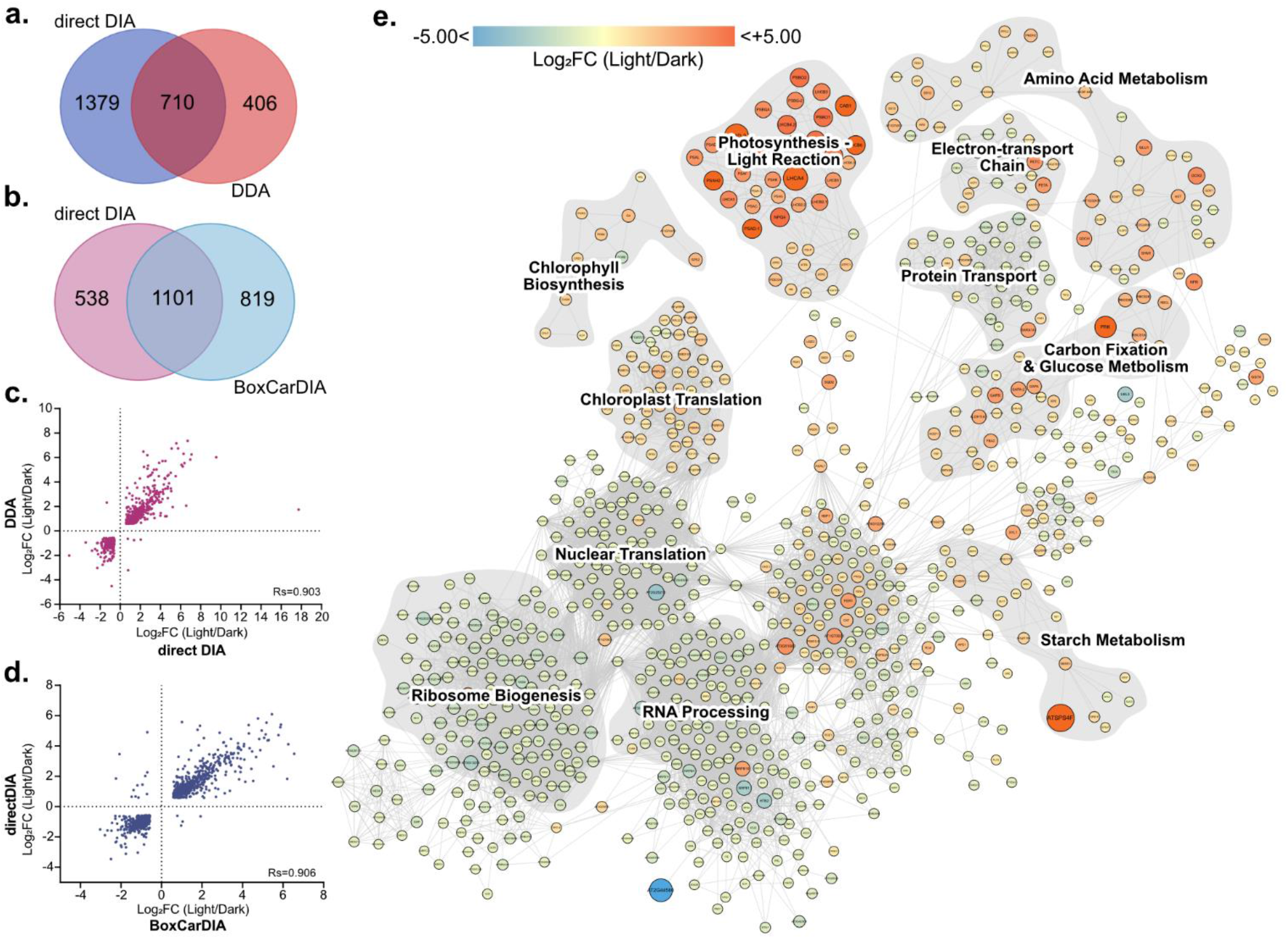
Differential protein abundance analysis for light- and dark-grown Arabidopsis cells. **(a.)** Venn diagram of protein groups with significantly changing protein abundances (q<0.05; Abs Log_2_FC>1.5) as measured by direct DIA and DDA. **(b.)** Venn diagram of protein groups with significantly changing protein abundances (q<0.05; Abs Log_2_FC>1.5) as measured by direct DIA and BoxCarDIA. **(c.)** Scatter plot of significant changes in protein abundance changes based on DDA and directDIA analysis. **(d.)** Scatter plot of significant changes in protein abundance changes based on directDIA and BoxCarDIA analysis. **(e.)** Association network of significantly changing proteins detected across all experiments. Network was constructed based on StringDB database and experiment datasets with a probability cut-off of 0.9. Only nodes with >3 edges are depicted. Clusters were manually annotated based on GO-terms and KEGG/Reactome pathway membership. Node sizes and color are scaled based on the median Log_2_FC (Light/Dark) from all analyses.

The Log_2_ Fold-Change values of proteins changing in both directDIA and DDA experiments were found to correlate to a high degree (Spearman’s R=0.9003), with proteins that were up-regulated in light- vs. dark-grown cells in directDIA analysis also up-regulated in DDA, and vice-versa (**Figure 7c**). A similar correlation was found between the Log_2_ Fold-Change values of proteins changing in both directDIA and BoxCarDIA analysis (**Figure 7d)**.

The complete dataset of 3,463 proteins changing significantly in abundance in light-vs dark-grown Arabidopsis cells is a valuable resource for future biochemical studies aiming to use these cell culture systems for protein interactomics experiments and other targeted proteomics analyses (**Supplementary Table 20**). To visualize the data, we further created a functional association network of these proteins by probing previously characterized databases and experiments compiled by StringDB^33^. This network validates our analysis, showing that clusters of proteins involved in photosynthesis, carbon-fixation, starch metabolism and amino-acid metabolism have increased abundance in light-vs. dark-grown cells, as expected (**Figure 7e**). Interestingly, clusters representing RNA processing, ER-Golgi transport, ribosome biogenesis, and nuclear translation are all downregulated, while chloroplast translation is upregulated, in light-vs. dark-grown cells (**Figure 7e**). These findings clearly highlight that the choice of cell culture growth condition is critical in order to avoid erroneous false positive and negative findings in protein interactomic experimentation.

## Conclusions

Until recently, DDA LC-MS (using both label-based and label-free approaches) has been the primary method of choice for proteomics studies in plants, due to the disadvantages of conventional DIA analysis, such as the requirement for project-specific spectral libraries. Here, we conclusively demonstrate that the newly developed library-free DIA proteomics approaches are vastly superior for plant proteomics as compared to currently used DDA methodologies. In particular, we find that our novel library-free BoxCarDIA method substantially improves upon gains provided by directDIA, and in doing so, solves the missing value problem in label-free proteomics. The advantages offered by BoxCarDIA includes: a greater number of protein identifications, greater dynamic range, and more robust protein quantification than DDA, with no change in instrumentation or increase in instrument analysis time. Our results, even using an advanced Tribrid Orbitrap-linear ion trap device, show that DDA acquisition is particularly inconsistent in its quantification of low-abundant proteins across samples. Similar results have been reported when comparing the abilities of directDIA and DDA to profile the phosphoproteome (a protein fraction with high dynamic range) of human tissue and cells ^20^. Our finding that more than 20% of identified proteins in a DDA experiment are detected in only 1 of 4 replicate injections of the same digest, and that these poorly quantified proteins tend to reside in the lower quartile of protein abundance, suggests an inherent drawback in DDA that has likely limited previous studies using this approach.

The data analyses undertaken here provide a useful template for benchmarking future quantitative mass spectrometry proteomics technologies from an end-user perspective. While our BoxCarDIA results demonstrate that segmented MS^1^ analysis through the use of BoxCar windows results in a variety of gains, there are likely further improvements in BoxCarDIA that may be realised through the use of better signal processing methods in order to reduce cycle times^34,35^. Our results argue persuasively for the widespread adoption of library-free BoxCarDIA for quantitative LFQ proteomics in plants. The demonstrated benefits in reproducibility and dynamic range of BoxCarDIA will be especially powerful for plant biology studies moving forward. In particular, proteomic analysis of multiple treatments (e.g., plant nutrition or herbicide studies), genotypes (e.g., breeding and selection trials), or timepoints (e.g., chronobiology studies), where comprehensive quantitative proteomic data are critical for maximizing our systems-level understanding of plants.

## Methods

### Arabidopsis cell culture

Heterotrophic *Arabidopsis thaliana*, cv. Ler suspension cells were obtained from the Arabidopsis Biological Resource Center (ABRC) and maintained in standard Murashige-Skoog media basal salt mixture (M524; PhytoTech Laboratories) at 21 °C as previously described^36^ under constant light (100 μmol m^-2^s^-1^) or constant dark. For the generation of experimental samples, 10 mL aliquots of each cell suspension (7 days old) were used to inoculate 8 separate 500 mL flasks that each contained 100 mL of fresh media. Experimental samples were grown for an additional 5 days prior to harvesting. Cells were harvested by vacuum filtration and stored at −80 °C.

### Sample Preparation

Quick-frozen cells were ground to a fine powder under liquid N2 using a mortar and pestle. Ground samples were aliquoted into 400 mg fractions. Aliquoted samples were then extracted at a 1:2 (w/v) ratio with a solution of 50 mM HEPES-KOH pH 8.0, 50 mM NaCl, and 4% (w/v) SDS. Samples were then vortexed and placed in a 95°C table-top shaking incubator (Eppendorf) at 1100 RPM for 15 mins, followed by an additional 15 mins shaking at room temperature. All samples were then spun at 20,000 x g for 5 min to clarify extractions, with the supernatant retained in fresh 1.5 mL Eppendorf tubes. Sample protein concentrations were measured by bicinchoninic acid (BCA) assay (23225; ThermoScientific). Samples were then reduced with 10 mM dithiothreitol (DTT) at 95°C for 5 mins, cooled, then alkylated with 30 mM iodoacetamide (IA) for 30 min in the dark without shaking at room temperature. Subsequently, 10 mM DTT was added to each sample, followed by a quick vortex, and incubation for 10 min at room temperature without shaking.

Total proteome peptide pools were generated using a KingFisher Duo (ThermoScientific) automated sample preparation device as outlined by Leutert et al. (2019)^37^ without deviation. Sample digestion was performed using sequencing grade trypsin (V5113; Promega), with generated peptide pools quantified by Nanodrop, acidified with formic acid to a final concentration of 5% (v/v) and then dried by vacuum centrifugation. Peptides were then dissolved in 3% ACN/0.1% TFA, desalted using ZipTip C18 pipette tips (ZTC18S960; Millipore) as previously described^7^, then dried and dissolved in 3.0% ACN/0.1% FA prior to MS analysis.

HeLa proteome analysis was carried out using a HeLa Protein Digest Standard (88329; Pierce). Four replicate injections of this digest per analysis type were carried out with the same methods as for Arabidopsis cell samples.

### Nanoflow LC-MS/MS analysis

Peptide samples were analysed using a Fusion Lumos Tribrid Orbitrap mass spectrometer (Thermo Scientific) in data dependent acquisition (DDA) and data independent acquisition (DIA) modes. Dissolved peptides (1 μg) were injected using an Easy-nLC 1200 system (LC140; ThermoScientific) and separated on a 50 cm Easy-Spray PepMap C18 Column (ES803A; ThermoScientific). The column was equilibrated with 100% solvent A (0.1% formic acid (FA) in water). Common MS settings between DDA and DIA runs included a spray voltage of 2.2 kV, funnel RF level of 40 and heated capillary at 300°C. All data were acquired in profile mode using positive polarity with peptide match off and isotope exclusion selected. All gradients were run at 300 nL/min with analytical column temperature set to 50°C.

#### DDA acquisition

Peptides were eluted with a solvent B gradient (0.1% (v/v) FA in 80% (v/v) ACN): 4% - 41% B (0 – 120 min); 41% - 98% B (120-125 min). DDA acquisition was performed using the Universal Method (ThermoScientific). Full scan MS^1^ spectra (350 - 2000 m/z) were acquired with a resolution of 120,000 at 200m/z with a normalized AGC Target of 125% and a maximum injection time of 50 ms. DDA MS^2^ were acquired in the linear ion trap using quadrupole isolation in a window of 2.5 m/z. Selected ions were HCD fragmented with 35% fragmentation energy, with the ion trap run in rapid scan mode with an AGC target of 200% and a maximum injection time of 100 ms. Precursor ions with a charge state of +2 - +7 and a signal intensity of at least 5.0e^3^ were selected for fragmentation. All precursor signals selected for MS/MS were dynamically excluded for 30s.

#### DIA acquisition

Peptides were eluted using a segmented solvent B gradient of 0.1% (v/v) FA in 80% (v/v) ACN from 4% - 41% B (0 - 107 min). DIA acquisition was performed as per Bekker-Jensen et al. (2020)^20^ and Biognosys AG. Full scan MS^1^ spectra (350 - 1400 m/z) were acquired with a resolution of 120,000 at 200 m/z with a normalized AGC Target of 250% and a maximum injection time of 45 ms. ACG target value for fragment spectra was set to 2000%. Twenty-eight 38.5 m/z windows were used with an overlap of 1 m/z **(Supplementary Table 21)**. Resolution was set to 30,000 using a dynamic maximum injection time and a minimum number of desired points across each peak set to 6.

BoxCar DIA acquisition was performed using the same gradient settings as DIA acquisition outlined above. MS^1^ analysis was performed by using two multiplexed targeted SIM scans of 10 BoxCar windows each. Detection was performed at 120,000 and normalized AGC targets of 100% per BoxCar isolation window. Isolation windows used are described in Supplementary Table 22. Windows were custom designed using the provided boxcarmaker R script that divides the MS1 spectra list into 20 m/z bins, each with an equal number of precursors, using the equal_freq function in the funModeling package (http://pablo14.github.io/funModeling/). Box sizes were scaled using this script applied to results from one of the directDIA runs.

MS^2^ acquisition was performed according to the settings described above for DIA acquisition.

### Raw data processing

DDA files were processed using MaxQuant software version 1.6.14^29,30^. MS/MS spectra were searched with the Andromeda search engine against a custom made decoyed (reversed) version of the Arabidopsis protein database from Araport 11^38^ concatenated with a collection of 261 known mass spectrometry contaminants. Trypsin specificity was set to two missed cleavage and a protein and PSM false discovery rate of 1%; respectively. Minimal peptide length was set to seven and match between runs option enabled. Fixed modifications included carbamidomethylation of cysteine residues, while variable modifications included methionine oxidation.

DIA files were processed with the Spectronaut directDIA experimental analysis workflow using default settings without N-acetyl variable modification enabled. Trypsin specificity was set to two missed cleavages and a protein and PSM false discovery rate of 1%; respectively. Data filtering was set to qQ-value (0.01) and global normalization with quantification performed at the MS2 level. For comparing BoxCarDIA and directDIA, the Spectronaut directDIA workflow was used with factory settings.

For hybrid (library- and library-free) DIA analysis, DDA raw files were first searched with the Pulsar search engine implemented in Spectronaut 14 to produce a search archive. Next, the DIA files were searched along with this search archive to generate a spectral library. The spectral library was then used for normal DIA analysis in Spectronaut 14. Default settings (without N-acetyl variable modification) were used in all steps. Final optimized Excalibur method files for DDA, directDIA and BoxCarDIA are provided as Supplemental Information.

### Data analysis

Downstream data analysis for DDA samples was performed using Perseus version 1.6.14.0^39^. Reverse hits and contaminants were removed, the data log_2_-transformed, followed by a data sub-selection criterion of n=3 of 4 replicates in at least one sample. Missing values were replaced using the normal distribution imputation method with default settings to generate a list of reliably quantified proteins. Subsequently, significantly changing differentially abundant proteins were determined and corrected for multiple comparisons (Bonferroni-corrected p-value < 0.05; q-value).

DirectDIA and BoxCarDIA data analysis was performed on Spectronaut v.14 using default settings.

Final numbers of PSMs, peptides, and protein groups identified were obtained from MaxQuant “summary.txt” files and from the result summary in Spectronaut.

Statistical analysis and plotting were performed using GraphPad Prism 8. Network analysis was performed on Cytoscape v.3.8.0 using the StringDB plugin.

### Data availability

Raw data have been deposited to the ProteomeExchange Consortium (http://proteomecentral.proteomexchange.org) via the PRIDE partner repository with the dataset identifier PXD022448. Source data used to produce all graphs is provided in the Supplemental Materials. R scripts and input data used can be downloaded from: https://github.com/UhrigLab/BoxCarMaker under a GNU Affero General Public License 3.0.

## Acknowledgements

The authors thank Jack Moore (University of Alberta) for assistance with operating the mass-spectrometer. We are grateful to Fabia Simona and Oliver Bernhardt (Biognosys AG) for assistance troubleshooting the Spectronaut software analysis, and to Florian Meier (Max Planck Institute for Biochemistry) for advice on BoxCar acquisition.

## Author Information

### Affiliations

**Department of Biological Sciences, University of Alberta, Edmonton T6G 2E9, Alberta, Canada**

Devang Mehta, Sabine Scandola, R. Glen Uhrig

### Contributions

D.M., and R.G.U contributed to Conceptualization, Methodology, and Formal Analysis. D.M. and S.S. contributed to Investigation. D.M. contributed to Visualization and Writing (original draft). R.G.U. performed Supervision and Funding Acquisition. D.M., S.S., and R.G.U contributed to Writing (review & editing).

### Corresponding author

Dr. R. Glen Uhrig

## Supplementary Tables

**Supplementary Table 1:** DDA protein quantification results for Arabidopsis cells

**Supplementary Table 2:** directDIA protein quantification results for Arabidopsis cells

**Supplementary Table 3:** Comparison of protein quantification for Arabidopsis cells between DDA and directDIA

**Supplementary Table 4:** DDA protein quantification results for HeLa digests

**Supplementary Table 5:** directDIA protein quantification results for HeLa digests

**Supplementary Table 6:** BoxCarDIA protein quantification results for Arabidopsis cells

**Supplementary Table 7:** directDIA protein quantification results for Arabidopsis cells in a second experiment for comparison with directDIA

**Supplementary Table 8:** Proteins identified in Arabidopsis cells using DDA

**Supplementary Table 9:** directDIA protein quantification results for Arabidopsis cells filtered for valid values in 3 of 4 replicates.

**Supplementary Table 10:** DDA protein quantification results for Arabidopsis cells with no imputation

**Supplementary Table 11:** directDIA protein quantification results for Arabidopsis cells filtered for valid values in all replicates.

**Supplementary Table 12:** DDA protein quantification results for Arabidopsis cells filtered for valid values in all replicates.

**Supplementary Table 13:** Proteins identified in HeLa digests using DDA

**Supplementary Table 14:** directDIA protein quantification results for HeLa digests filtered for valid values in 3 of 4 replicates.

**Supplementary Table 15:** DDA protein quantification results for HeLa digests with no imputation

**Supplementary Table 16:** directDIA protein quantification results for HeLa digests filtered for valid values in all replicates.

**Supplementary Table 17:** DDA protein quantification results for HeLa digests filtered for valid values in all replicates.

**Supplementary Table 18:** BoxCarDIA protein quantification results for HeLa digests

**Supplementary Table 19:** directDIA protein quantification results for HeLa digests in a second experiment for comparison with BoxCarDIA

**Supplementary Table 20:** Proteins changing significantly in abundance between light and dark-grown Arabidopsis cells, measured using both directDIA and DDA.

**Supplementary Table 21:** Precursor selection mass list table

**Supplementary Table 22:** BoxCar isolation windows

## Supplementary Figures

**Figure S1:**
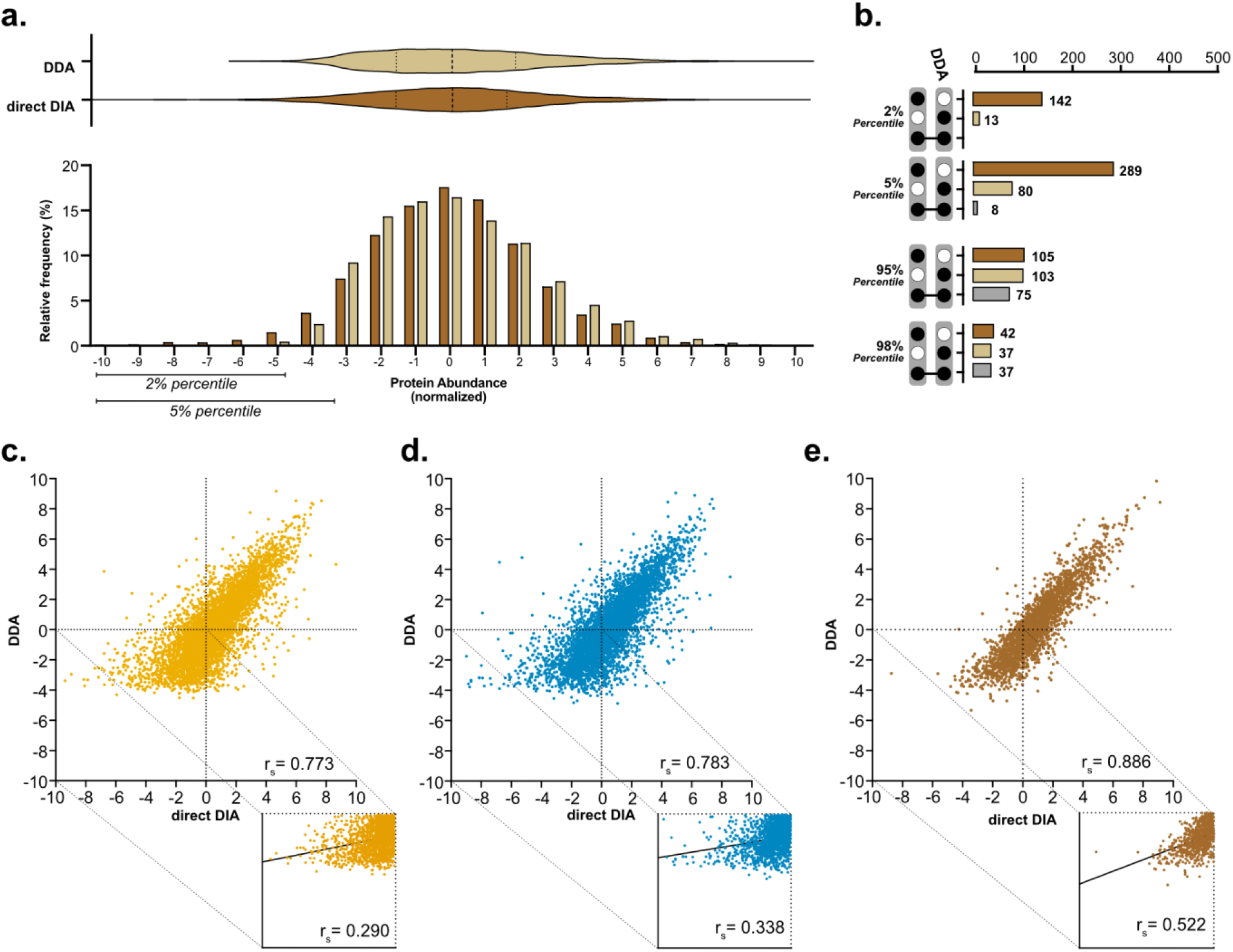
Comparison of protein quantification results using DDA and directDIA analysis. **(a.)** Frequency distribution of normalized protein abundances for DDA and directDIA analysis and corresponding violin plots with median and quartile lines marked for HeLa digests. **(b.)** Upset plots depicting intersections in protein groups quantified by DDA and direct DIA at either extreme of the abundance distribution for HeLa digests. **(c.-e.)** Scatter plots of protein groups quantified by DDA and direct DIA for light-grown Arabidopsis cells, dark-grown Arabidopsis cells, and HeLa digests. Insets show correlations for protein groups with abundances less than the median.

**Figure S2:**
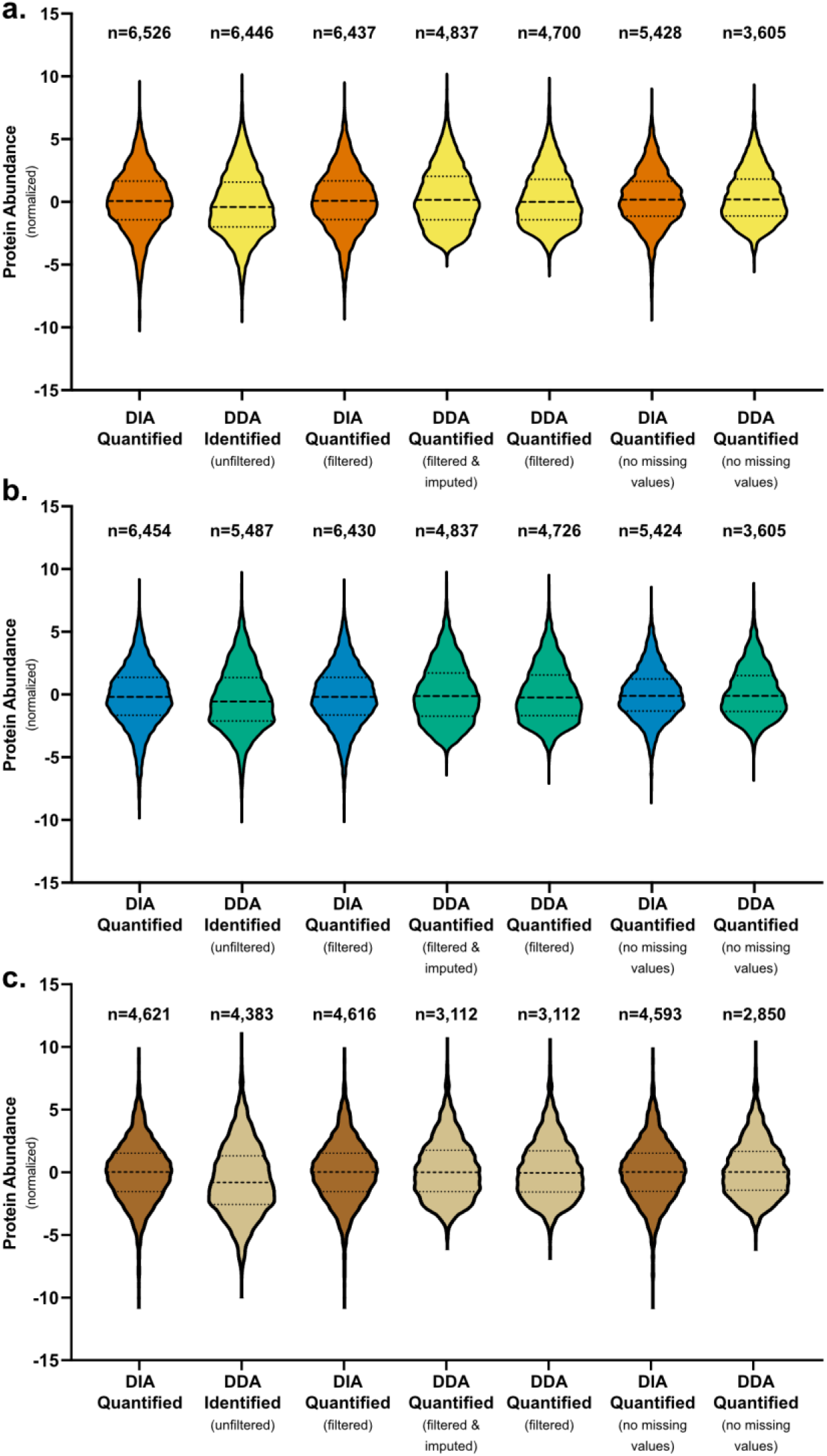
Protein abundance distributions by analysis type and data filtering settings. Violin plots showing normalized protein abundance for proteins quantified by direct DIA (default setting), identified by DDA, quantified by DIA (filtered for protein groups present in at least 3 samples in any one condition), quantified by DDA (filtered for protein groups present in at least 3 samples in any one condition with missing values imputed), quantified by DDA (filtered for protein groups present in at least 3 samples in any one condition with missing values left blank), quantified by DIA (counting only protein groups found in all samples), and quantified by DIA (counting only protein groups found in all samples), respectively for **(a.)** light grown Arabidopsis cells **(b.)** dark grown Arabidopsis cells and **(c.)** HeLa cell digestion standards. (n= number of protein groups).

**Figure S3:**
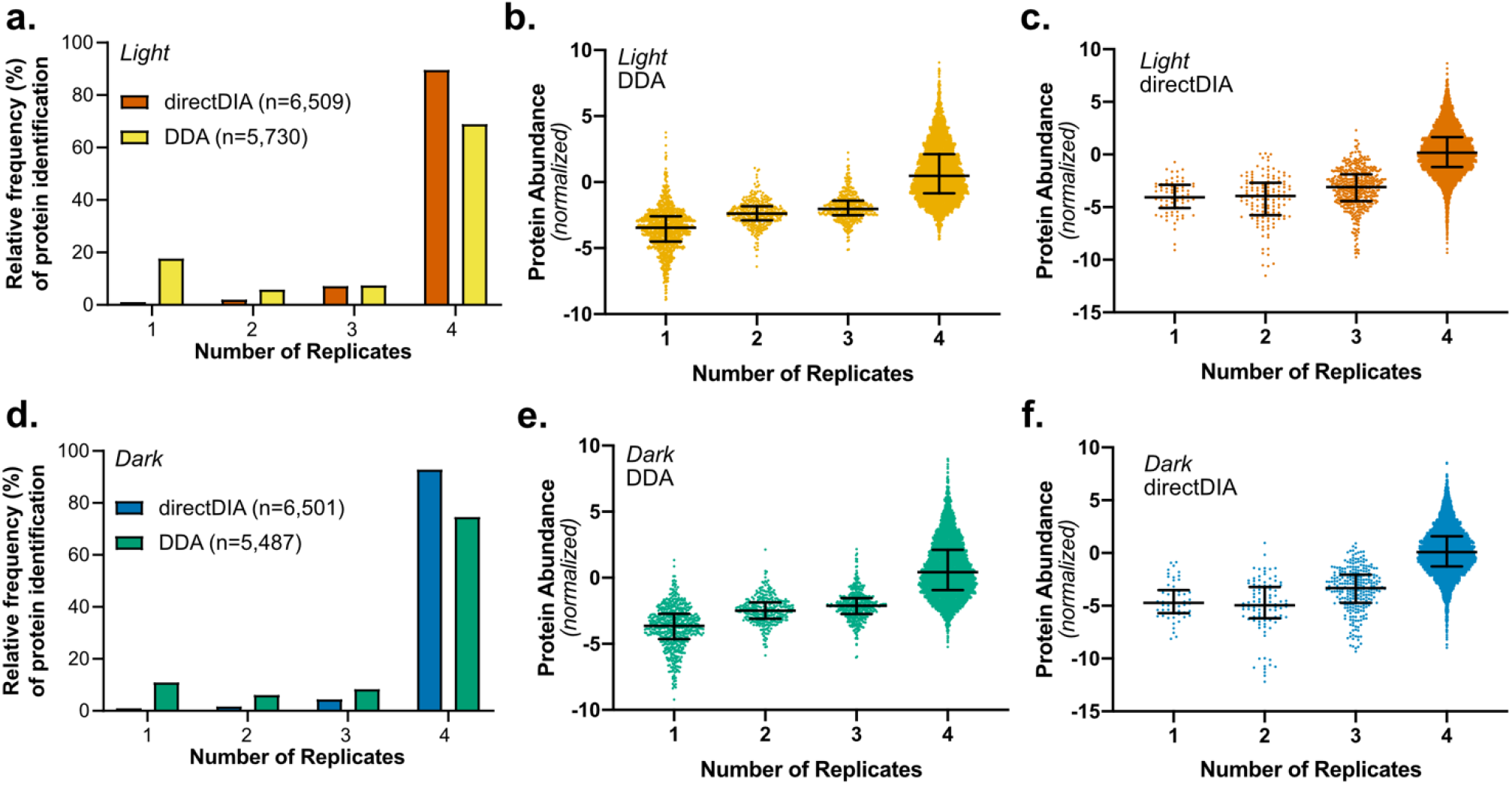
The DDA missing value problem explains the gap in quantification of low abundant proteins compared to direct DIA. **(a.)** Histograms of direct DIA or DDA protein group identifications across replicate samples for light-grown Arabidopsis cells. **(b & c)** Normalized abundances of proteins binned by the number of replicates containing each protein for direct DIA and DDA. Bars represent median and interquartile range. **(d.-f.)** Same as above for dark-grown cells.

**Figure S4:**
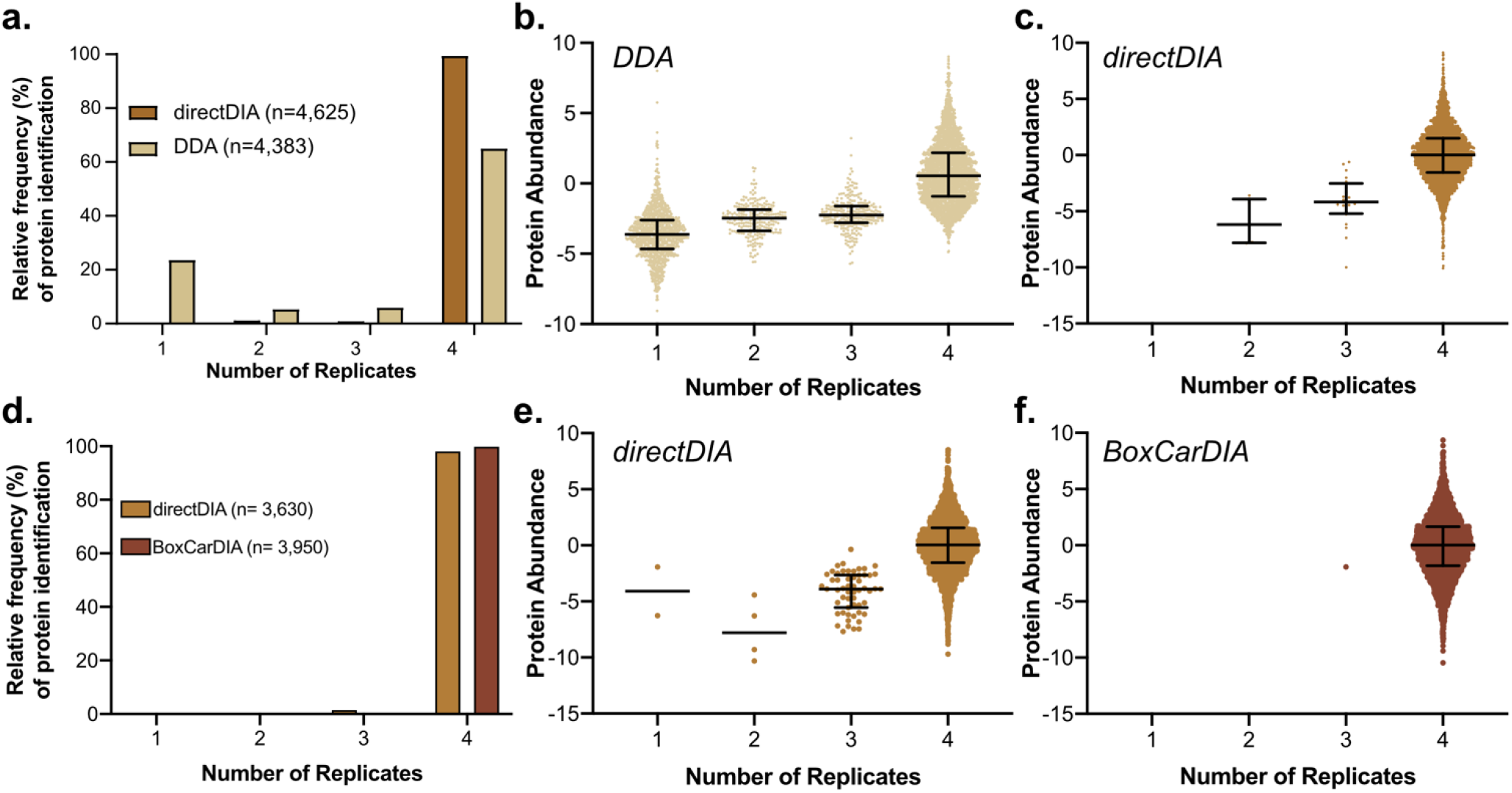
BoxCarDIA can quantify peptides and protein groups consistently between independent technical replicate injections of HeLa digests. **(a.)** Histograms of BoxCarDIA, directDIA, or DDA peptide identifications across replicate injections of HeLa digests. **(b-d.)** Normalized abundances of peptides binned by the number of replicates containing each protein for DDA, directDIA and BoxCarDIA. Bars represent median and interquartile range. **(e-h)** Same as above for protein group identifications.

**Figure S5:**
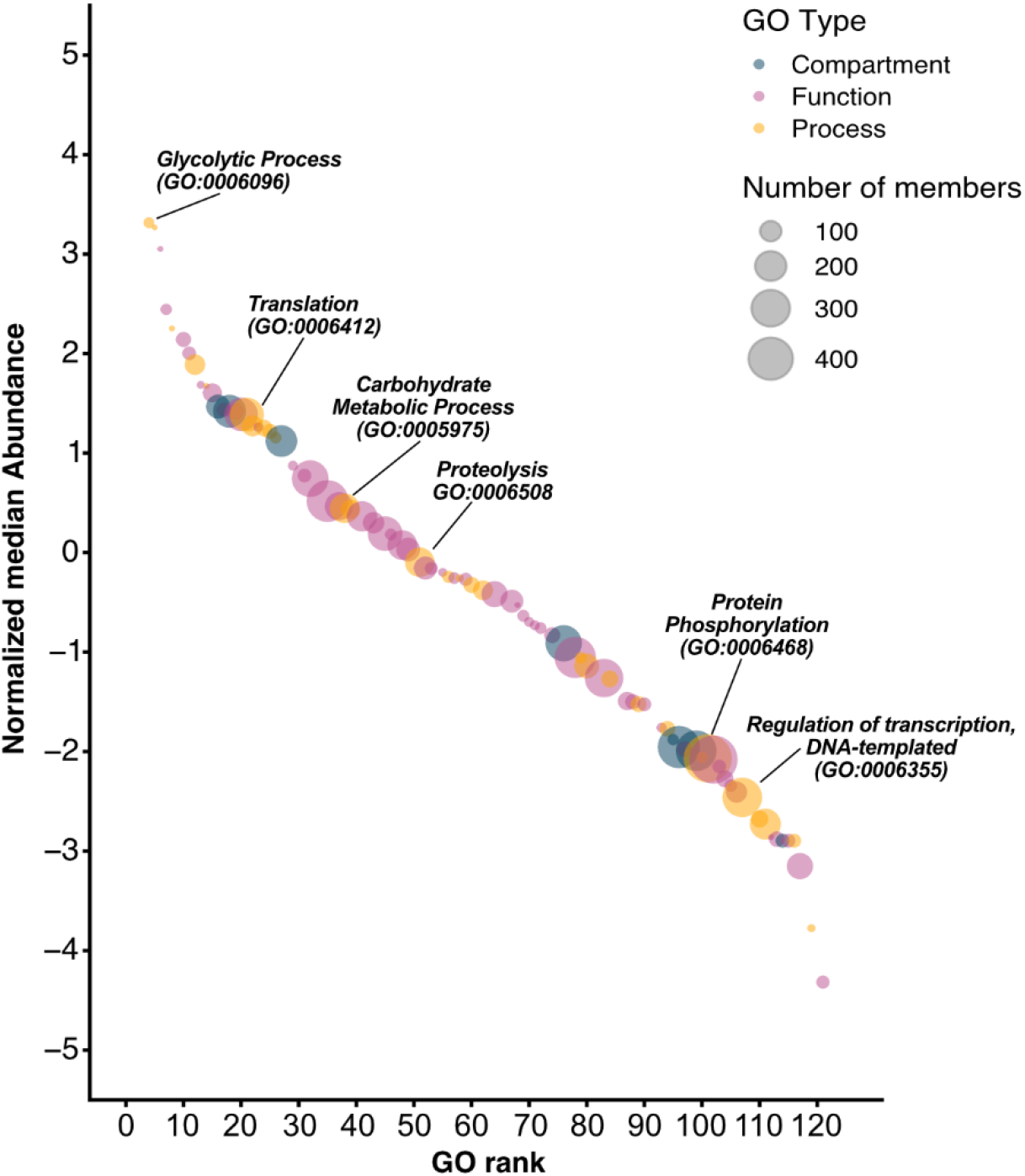
Arabidopsis Gene Ontology categories ranked by the abundance of their constituent proteins. Data from a deep proteome analysis of Arabidopsis cells performed by Mergner et al., 2020.

## References

1. Clark, N. M. et al. Integrated omics networks reveal the temporal signaling events of brassinosteroid response in Arabidopsis. BioRxiv (2020). doi:10.1101/2020.09.04.283788

2. Mehta, D. et al. Phosphate and phosphite have a differential impact on the proteome and phosphoproteome of Arabidopsis suspension cell cultures. Plant J. 105, 924–941 (2021).

3. Vanderschuren, H. et al. Large-Scale Proteomics of the Cassava Storage Root and Identification of a Target Gene to Reduce Postharvest Deterioration. Plant Cell 26, 1913–1924 (2014).

4. Ting, L., Rad, R., Gygi, S. P. & Haas, W. MS3 eliminates ratio distortion in isobaric multiplexed quantitative proteomics. Nat. Methods 8, 937–940 (2011).

5. Ow, S. Y. et al. iTRAQ underestimation in simple and complex mixtures: “the good, the bad and the ugly”. J. Proteome Res. 8, 5347–5355 (2009).

6. Graf, A. et al. Parallel analysis of Arabidopsis circadian clock mutants reveals different scales of transcriptome and proteome regulation. Open Biol 7, (2017).

7. Uhrig, R. G., Schläpfer, P., Roschitzki, B., Hirsch-Hoffmann, M. & Gruissem, W. Diurnal changes in concerted plant protein phosphorylation and acetylation in Arabidopsis organs and seedlings. Plant J. 99, 176–194 (2019).

8. Hartl, M. et al. Lysine acetylome profiling uncovers novel histone deacetylase substrate proteins in Arabidopsis. Mol. Syst. Biol. 13, 949 (2017).

9. Aebersold, R. & Mann, M. Mass-spectrometric exploration of proteome structure and function. Nature 537, 347–355 (2016).

10. Gillet, L. C. et al. Targeted data extraction of the MS/MS spectra generated by data-independent acquisition: a new concept for consistent and accurate proteome analysis. Mol. Cell Proteomics 11, O111.016717 (2012).

11. Ludwig, C. et al. Data-independent acquisition-based SWATH-MS for quantitative proteomics: a tutorial. Mol. Syst. Biol. 14, e8126 (2018).

12. Tsou, C.-C. et al. DIA-Umpire: comprehensive computational framework for data-independent acquisition proteomics. Nat. Methods 12, 258–64, 7 p following 264 (2015).

13. Li, Y. et al. Group-DIA: analyzing multiple data-independent acquisition mass spectrometry data files. Nat. Methods 12, 1105–1106 (2015).

14. Biognosys AG. A new era in proteomics: spectral library free data independent acquisition (directDIA). The Analytical Scientist (2017).

15. Bruderer, R., Bernhardt, O. M., Gandhi, T. & Reiter, L. High-precision iRT prediction in the targeted analysis of data-independent acquisition and its impact on identification and quantitation. Proteomics 16, 2246–2256 (2016).

16. Reiter, L. MP 125: Direct Searching of DIA Data Catches up with Sample-specific Libraries. in Proceedings of the 68th ASMS Conference on Mass Spectrometry and Allied Topics, Online Meeting (American Society for Mass Spectrometry, 2020). at <https://biognosys.com/media.ashx/mp125lukasreiterasms2020.pdf>

17. Yang, Y. et al. In silico spectral libraries by deep learning facilitate data-independent acquisition proteomics. Nat. Commun. 11, 146 (2020).

18. Demichev, V., Messner, C. B., Vernardis, S. I., Lilley, K. S. & Ralser, M. DIA-NN: neural networks and interference correction enable deep proteome coverage in high throughput. Nat. Methods 17, 41–44 (2020).

19. Muntel, J. et al. Surpassing 10 000 identified and quantified proteins in a single run by optimizing current LC-MS instrumentation and data analysis strategy. Mol. Omics 15, 348–360 (2019).

20. Bekker-Jensen, D. B. et al. Rapid and site-specific deep phosphoproteome profiling by data-independent acquisition without the need for spectral libraries. Nat. Commun. 11, 787 (2020).

21. Meier, F., Geyer, P. E., Virreira Winter, S., Cox, J. & Mann, M. BoxCar acquisition method enables single-shot proteomics at a depth of 10,000 proteins in 100 minutes. Nat. Methods 15, 440–448 (2018).

22. Van Leene, J. et al. Capturing the phosphorylation and protein interaction landscape of the plant TOR kinase. Nat. Plants 5, 316–327 (2019).

23. Van Leene, J. et al. Targeted interactomics reveals a complex core cell cycle machinery in Arabidopsis thaliana. Mol. Syst. Biol. 6, 397 (2010).

24. Gonzalez, N. et al. A repressor protein complex regulates leaf growth in arabidopsis. Plant Cell 27, 2273–2287 (2015).

25. Antosz, W. et al. The Composition of the Arabidopsis RNA Polymerase II Transcript Elongation Complex Reveals the Interplay between Elongation and mRNA Processing Factors. Plant Cell 29, 854–870 (2017).

26. Dejonghe, W. et al. Disruption of endocytosis through chemical inhibition of clathrin heavy chain function. Nat. Chem. Biol. 15, 641–649 (2019).

27. Marondedze, C., Thomas, L., Serrano, N. L., Lilley, K. S. & Gehring, C. The RNA-binding protein repertoire of Arabidopsis thaliana. Sci. Rep. 6, 29766 (2016).

28. Arora, D. et al. Establishment of Proximity-dependent Biotinylation Approaches in Different Plant Model Systems. Plant Cell (2020). doi:10.1105/tpc.20.00235

29. Cox, J. & Mann, M. MaxQuant enables high peptide identification rates, individualized p.p.b.-range mass accuracies and proteome-wide protein quantification. Nat. Biotechnol. 26, 1367–1372 (2008).

30. Tyanova, S., Temu, T. & Cox, J. The MaxQuant computational platform for mass spectrometry-based shotgun proteomics. Nat. Protoc. 11, 2301–2319 (2016).

31. Lex, A., Gehlenborg, N., Strobelt, H., Vuillemot, R. & Pfister, H. Upset: visualization of intersecting sets. IEEE Trans Vis Comput Graph 20, 1983–1992 (2014).

32. Mergner, J. et al. Mass-spectrometry-based draft of the Arabidopsis proteome. Nature 579, 409–414 (2020).

33. Szklarczyk, D. et al. STRING v11: protein-protein association networks with increased coverage, supporting functional discovery in genome-wide experimental datasets. Nucleic Acids Res. 47, D607–D613 (2019).

34. Grinfeld, D., Aizikov, K., Kreutzmann, A., Damoc, E. & Makarov, A. Phase-Constrained Spectrum Deconvolution for Fourier Transform Mass Spectrometry. Anal. Chem. 89, 1202–1211 (2017).

35. Meier, F. High Dynamic Range Proteome Analysis with BoxCar DIA and Super-Resolution Orbitrap Mass Spectrometry. (2020). at <http://assets.thermofisher.com/TFS-Assets/CMD/posters/po-65792-ms-proteome-boxcar-dia-orbitrap-asms2020-po65792-en.pdf>

36. Uhrig, R. G. & Moorhead, G. B. Two ancient bacterial-like PPP family phosphatases from Arabidopsis are highly conserved plant proteins that possess unique properties. Plant Physiol. 157, 1778–1792 (2011).

37. Leutert, M., Rodríguez-Mias, R. A., Fukuda, N. K. & Villén, J. R2-P2 rapid-robotic phosphoproteomics enables multidimensional cell signaling studies. Mol. Syst. Biol. 15, e9021 (2019).

38. Cheng, C.-Y. et al. Araport11: a complete reannotation of the Arabidopsis thaliana reference genome. Plant J. 89, 789–804 (2017).

39. Tyanova, S. et al. The Perseus computational platform for comprehensive analysis of (prote)omics data. Nat. Methods 13, 731–740 (2016).

